# Carbon Sequestration Potential of Oil Palm Plantations in Southern Philippines

**DOI:** 10.1101/2020.04.14.041822

**Authors:** Sheila Mae C. Borbon, Michael Arieh P. Medina, Jose Hermis P. Patricio, Angela Grace Toledo-Bruno

**Affiliations:** Department of Environmental Science, College of Forestry and Environmental Science Central Mindanao University, University Town, Musuan, Bukidnon, Philippines

**Keywords:** carbon stock assessment, oil palm, biomass, carbon sequestration, carbon pools

## Abstract

Aside from the greenhouse gas reduction ability of palm oil-based biofuel as alternative to fossil fuels, another essential greenhouse gas mitigation ability of oil palm plantation is in terms of offsetting anthropogenic carbon emissions through carbon sequestration. In this context, this study was done to determine the carbon sequestration potential of oil palm plantations specifically in two areas in Mindanao, Philippines. Allometric equation was used in calculating the biomass of oil palm trunk. Furthermore, destructive methods were used to determine the biomass in other oil palm parts (fronds, leaves, and fruits). Carbon stocks from the other carbon pools in the oil palm plantations were measured which includes understory, litterfall, and soil. Results revealed that the average carbon stock in the oil palm plantations is 40.33 tC/ha. Majority of the carbon stock is found in the oil palm plant (53%), followed by soil (38%), litterfall (6%), and understory, (4%). The average carbon sequestration rate of oil palm plants is estimated to be 4.55 tC/ha/year. It is advised that oil palm expansions should be done on grasslands and barren lands instead of forest lands to avoid incurring “carbon debts”. It is also recommended that oil palms should be mixed with shrub crops species to enhance soil organic carbon as well as increase the aboveground biomass in oil palm plantations.

## 1. Introduction

The world’s remaining forests are currently under threat due to the ongoing massive expansion of oil palm plantations (Koh and Wilcove 2007). Oil palm is preferable because it has a higher oil yield per unit area compared to other oil crops. It produces two types of oils from the same fruit – palm oil from the flesh or mesocarp and palm kernel oil from the seed or kernel inside the hard-shell mesocarp. Aside from that, its kernels also produce a residual product known as palm kernel meal, a raw material for animal feed (Basiron 2007). With the ubiquitous utilization of palm oil globally, the oil palm industry became one of the major sources of raw material for food, medicinal, chemical and industrial uses (Sanquetta et al 2015).

The Philippines has been cultivating and processing palm oil for the past three decades although this industry is considered small compared to the millions of hectares of oil palm plantations in Malaysia and Indonesia. Nevertheless, the continuing demand for palm oil both in the local and global market provided an avenue for accelerating the expansion of oil palm plantation in the country (Villanueva 2011). In addition, the expansion of oil palm plantations in the country was also encouraged due to the enactment of the Biofuels Act of 2011, a law that mandates a 2% biodiesel blend for diesel and 10% bioethanol blend in gasoline (Pulhin et al 2014).

Based on report from the Philippine Palm Oil Development Council (PPODC), 46,608 hectares have been planted with oil palm in the Philippines in 2009. This constitutes a 160% increase of oil palm plantation area since 2005. Most of these areas are found in Mindanao while a few can be found in Palawan and Central Visayas (Villanueva 2011).

In the context of climate change mitigation, the use of palm oil for biofuels constitutes a substantial greenhouse gas reduction since biofuels have lesser carbon emissions than fossil fuels. Furthermore, another climate change mitigation potential of oil palm is the offsetting of anthropogenic carbon emissions through its carbon storage ability (Lamade and Bouillet 2005, Morel et al 2011, Pulhin et al 2014, Sanquetta et al 2015).

Putting into consideration the disastrous effect of oil palm expansion to biodiversity (referring to forest conversion into oil palm plantations) (Koh and Wilcove 2007), this study deals with the potential of oil palm plantations in avoiding “carbon debts” (referring to conversion of grassland or barren lands and not forestlands) (Frazão et al 2012, Sanquetta et al 2015). Previous studies have been done with regards to the carbon stock of oil palm plantations in other countries (Thenkabail et al 2004, Syahrinudin 2005, Lamade and Bouillet 2005, Morel et al 2011, Sanquetta et al 2015, Yuliyanto 2016). In the Philippines, Pulhin et al (2014) studied the carbon storage of oil palm parts (trunk, fronds, leaves, fruit, flowers, and roots) which provided a relevant baseline on the carbon stock potential of oil palm in the context of the Philippines. However, this study aims to augment the said study by including other carbon pools in oil palm plantations (understory, litterfall, and soil) aside from the oil palm parts. This hopefully will provide comprehensive information regarding oil palm carbon stock in the country at the plantation level.

## 2. Materials and Methods

### 2.1 Location of the Study

Two oil palm plantations were considered for the study. The first plantation is located in the Municipality of Kadingilan, Bukidnon Province specifically in Cabadiangan Village. The second plantation is located in the Municipality of Montevista, in the Province of Compostela Valley specifically in Bankeruhan Sur Village. Both of these areas are located in the southern island of Mindanao, Philippines (Fig. 1).

**Fig. 1.**
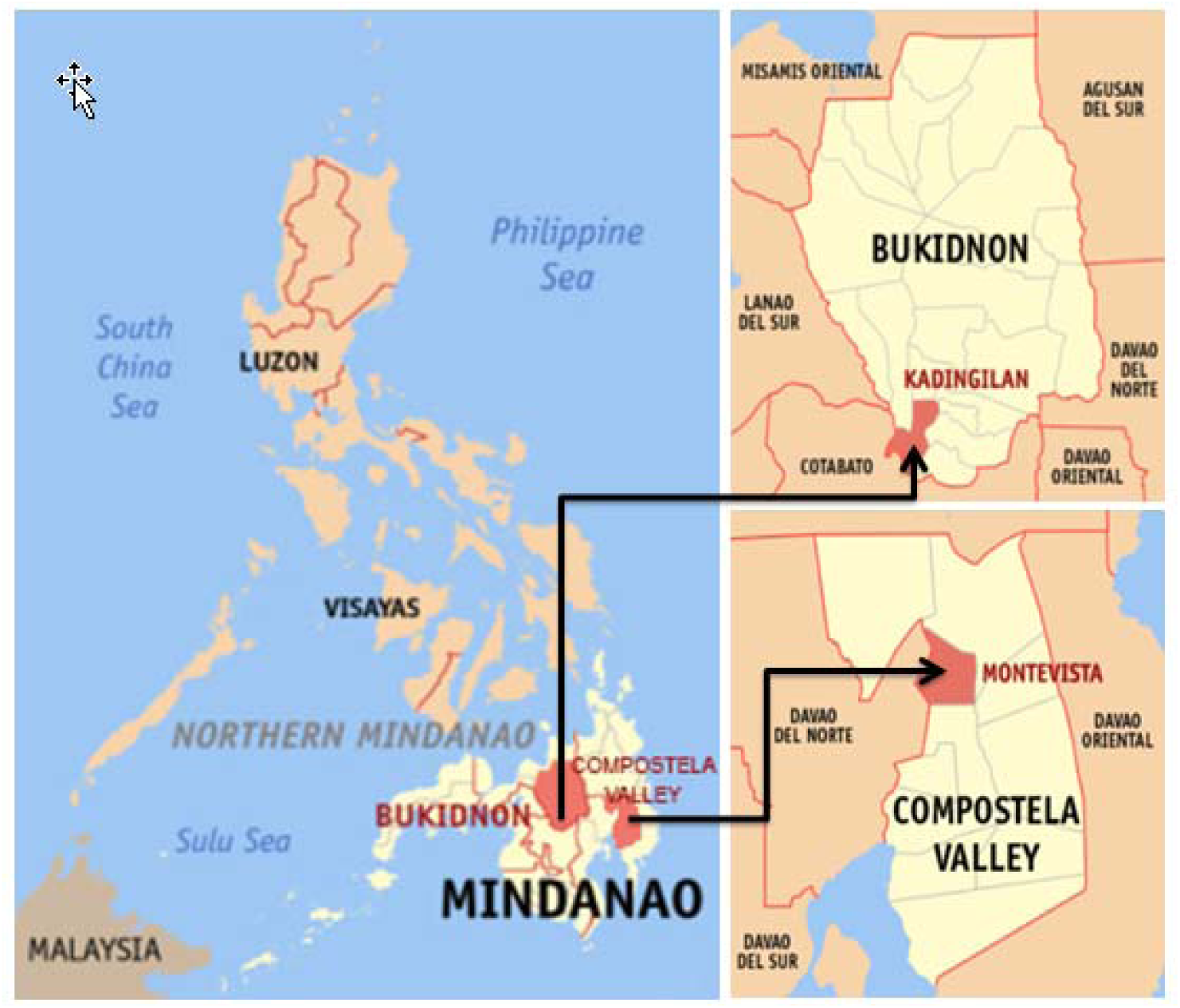
Location of the Study Area (*Source: WikiMedia Commons*)

The climatic condition of the Municipality of Kadingilan is considered as type C agro-climatic zone. This zone covers most potential agricultural areas in the province of Bukidnon. The heaviest rainfall in the area occurs during the months of May to September. The topography of the area is characterized as rugged terrain with rolling, hilly and mountainous portion. On the other hand, the climatic condition of the Municipality of Montevista falls under type IV, characterized by not very pronounced maximum rain period and no dry seasons. The topography of the area is classified into level to undulating, moderately sloping rolling, rolling to hills, steep hills and mountainous.

### 2.2 Sampling Procedure

Two 0.05 hectare circular plots (r =12.6 m, consisting of 7 oil palm) were randomly established in 2, 4, 6 and 7 year old oil palm plantations in Bankeruhan Sur, Montevista, Compostela Valley and 2, and 7 year old plantations in Barangay Cabadiangan, Kadingilan,Bukidnon. Thus, 8 plots were established in Bankeruhan Sur, Montevista, Compostela Valley (4 stand ages, 2 replications) and 4 plots in Kadingilan, Bukidnon (2 stand ages, 2 replicates). Details of this sampling procedure can be found in Syahrinudin (2005).

### 2.3 Biomass and Carbon Storage Calculation of Oil Palm

Biomass of oil palm trunk was calculated using the allometric equation specified by Morel et al (2012). The equation used to calculate this is as follows: trunk biomass is indicated as T_b_ which was calculated using Eq. (1). Furthermore, trunk density (ρ) was calculated using Eq. (2).

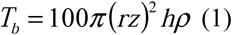

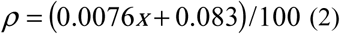

Where;

r = radius of the trunk (in cm) without frond bases

ρ = trunk density in kg/m^3^ dependent on the age

x = age of the palm in years

z = ratio of the diameter below frond bases, estimated to be 0.777 h= height of the trunk (in m) to the base of the frond

Other oil palm parts (fronds, leaves, and fruits) were sampled destructively. One frond was taken from one oil palm per plot which were chopped and placed in plastic bags and weighed. Leaves from that same frond were also taken as a separate sample using the same procedure above. Furthermore, fruits from one oil palm per plot were also gathered as samples.

A 1 kilogram sample from each oil palm parts (fronds, leaves, fruits) were then subjected to oven drying. However, prior to oven drying, the collected samples were air dried for one week to shorten oven drying time. The samples were oven dried at a temperature of 100°C for three days or until the weights of the samples became constant. Biomass value for the leaves, flowers, fruits, (ODWt) was calculated using Eq. (3) (Pulhin et al 2014):

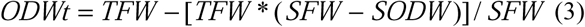

Where:

ODWt= total oven dry weight

TFW = total fresh weight

SFW = sample fresh weight

SODW = sample oven-dry weight

Total biomass of the oil palm was estimated by summing the biomass values derived for each of the plant parts: trunk, fronds, leaves, and fruits. Furthermore, carbon stored was determined by multiplying the total biomass for each plant parts with corresponding carbon content values. For trunk, a default value (40.64%) was used based on previous literature (UNEP, 2012). For the rest of oil palm parts (frond, leaves, and fruits), carbon content used was based on the results of the laboratory analysis of the samples brought to the Soil and Plant Analysis Laboratory in Central Mindanao University, Bukidnon.

### 2.4 Biomass and Carbon Storage Calculation of Understory and Litter

For understory, nine (9) 0.5m x 0.5m (0.25 sq.m.) sub plots were randomly established to assess the biomass of the herbaceous undergrowth occupying the respective plot. All undergrowth in these sub-plots were harvested and weighed to determine the total fresh weight. Likewise, in the same sub plots as understory, litterfall (i.e. any tree necromass<5cm in diameter and/or <30 cm length, undecomposed plant materials or crop residues, and all unburned leaves and branches) were also collected in the same sub plots as the understory.

Furthermore, 300g subsamples were subjected to air and oven drying. Oven drying was set at 65-70 ºC and observed for at least 48 hours or until the samples reached their stable weight. Oven-dry weights of subsamples were determined to compute for the total dry weights using Eq. (4) (Hairiah et al 2001):

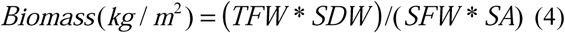

Where:

TFW = Total Fresh Weight (kg)

SDW = Subsample dry weight (g)

SFW = Subsample Fresh Weight (g)

SA = Sample Area (m^2^)

A small sample (2 grams) of each one of the understory and litterfall were analyzed for carbon content at the SPAL (Soil and Plant Analysis Laboratory). Carbon storage is then computed by multiplying understory and litterfall biomass with their respective carbon content.

### 2.5 Soil Organic Carbon

In each sampling site, soil samples were taken from 5-10 cm, 10-20 cm, and 20-30 soil depth. Then, a composite sample of about ½ kg was taken for soil analysis to the Soil and Plant Analysis Laboratory (SPAL) of Central Mindanao University to determine the Soil Organic Carbon (SOC) using the Walkey-Black Method.

Furthermore, undisturbed soil cores were collected using a sampling metal tube with a diameter of 5.3 cm and a length of 10 cm. In order to avoid soil loss from its cores, the sample was wrapped in an aluminum foil. Samples were brought to SPAL, for oven drying to constant weight for 40 hours at ± 105°C. Bulk density (BD) and Soil Organic Carbon (SOC) were computed using Eqs. (5) and (6), respectively (Patricio 2014):

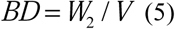

Where:

BD = bulk density of the soil sample

W_2_ = oven dry weight of the sample

V = volume of the cylinder/tube.

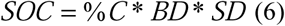

Where:

SOC = Soil Organic Carbon (MgC/ha)

BD = Bulk Density (Mg/m3)

SD = Sampling Depth (cm)

## 3. Results and Discussions

### 3.1 Biomass of Oil Palm Parts

As shown in Table 1, the average biomass of oil palms is around 52.57 tons/ha. This ranges from 21.84 – 53.03 tons/ha depending on the age of the plantation. Majority of the biomass is found in the trunk which comprises around 82% of the total biomass. Trunk biomass ranges from 15.49 tons/ha to 70.72 tons/ha with an average of 42.99 tons/ha. In a study by Asari et al (2013) of 60 oil palm stands in Malaysia, oil palm trunk biomass ranges from 8.53 to 70.94 tons/ha with an average of 33.84 tons/ha. The said oil palm stands ranges in age from 6 to 23 years old. In a similar study, trunks have been found to account for most of the total biomass in oil palm (Pulhin et al 2014). On the other hand, fronds and leaves comprise around 18% of the total biomass though it has been found to have higher percentage in previous literature. Fronds biomass in the study ranges from 3.18 tons/ha to 7.86 tons/ha averaging around 5.28 tons/ha. Leaf biomass ranges from 2.39 tons/ha to 5.61 tons/ha with an average of 4.15 tons/ha. In another study, frond biomass ranges from 4.50 tons/ha to 9.75 tons/ha or an average of 6.92 tons/ha (Asari et al 2013).

**Table 1.** Biomass of the Different Oil Palm Parts

| LOCATION | AGE<br>(yrs) | BIOMASS (tons/ha) |  |  |  |  |
| --- | --- | --- | --- | --- | --- | --- |
|  |  | TRUNK | FRONDS | LEAVES | FRUITS | TOTAL |
| Kadingilan, Bukidnon | 2 | 18.70 | 3.18 | 2.39 | 0.11 | 24.38 |
|  | 4 | 34.16 | 4.70 | 3.94 | 0.22 | 43.02 |
|  | 6 | 51.78 | 5.57 | 4.15 | 0.20 | 61.70 |
|  | 7 | 70.72 | 6.57 | 5.47 | 0.26 | 83.02 |
| Montevista, Compostela Valley | 2 | 15.49 | 3.27 | 3.00 | 0.08 | 21.84 |
|  | 7 | 68.78 | 7.86 | 5.61 | 0.14 | 82.39 |
| **Mean** |  | **42.99** | **5.285** | **4.15** | **0.155** | **52.57** |

Fruits make up less than 1% of the total biomass in this study which ranges from 0.08 tons/ha to 0.26 tons/ha with an average of 0.15 tons/ha. Previous studies have reported that fruits are found to have the lowest biomass percentage among the oil palm parts (Pulhin et al 2014).

### 3.2 Understory and Litter Biomass of the Oil Palm Plantations

As shown in Table 2 the mean biomass of the understory in the two oil palm plantations is 3.71 tons/ha. This result is comparable to the understory biomass of a 20-year-old palm plantation in Indonesia (3.6 tons/ha) (Syahrinudin 2005). In comparison, fruit tree plantations have a much lower understory biomass (0.78 – 1.51 tons/ha) compared to the study results (Janiola and Marin 2016). Same is true with understory biomass in agroforestry farms (0.16 to 2.29 tons/ha) (Labata et al 2012). However, forest tree plantations have extremely higher understory biomass ranging 8.5 tons/ha to 24.88 tons/ha (Tulod 2015).

**Table 2.**
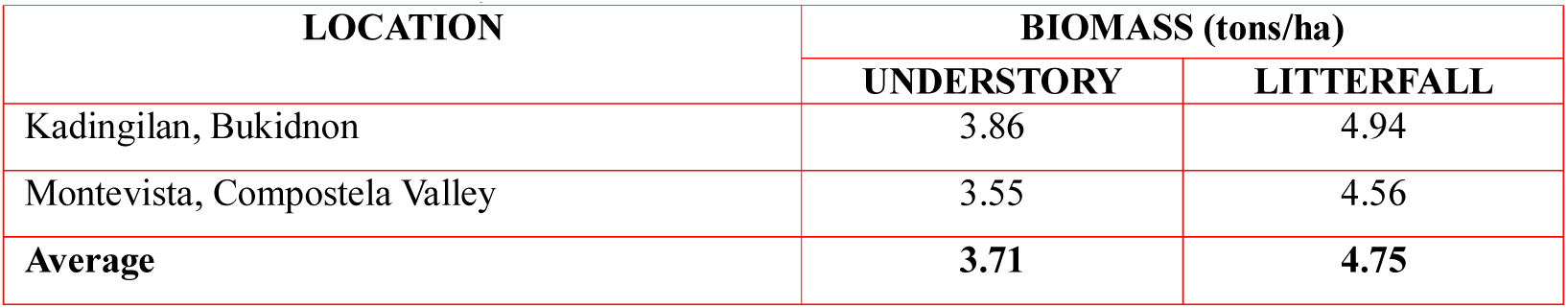
Biomass of Understory and Litterfall in the Oil Palm Plantations

On the other hand, the litterfall biomass had an average of 4.75 ton/ha in the study areas. Results from Syahrinudin (2005) have almost the same result with regards to litterfall biomass of a 3-year oil palm plantation (4.7 tons/ha). Furthermore, the above results fall within the observed range of litterfall biomass for fruit tree plantations (3.25 - 6.93 tons/ha) (Janiola and Marin 2016). Litterfall biomass in agroforestry farms however has a bit lower observed range (0.42 – 4.5 tons/ha) (Labata et al 2012). Comparably though, forest tree plantations have a higher observed range of litterfall biomass (8.96 – 42.83 tons/ha). Furthermore, bamboo plantations seem to have a higher litterfall biomass at 9.45 tons/ha (Toledo-Bruno et al 2017) than what was observed in the study area.

### 3.3 Carbon Storage in Oil Palm

As shown in Table 3, the average carbon stock of oil palm is 21.34 tC/ha ranging from 8.63 tC/ha to 33.76 tC/ha depending on the age of the plantation. A study by Pulhin et al (2014) on oil palms in Bohol, Philippines aged two, five, seven, and nine years has an average carbon stock of 9.6 tC/ha which is a bit similar to the results of the study. Previous studies abroad also has approximately similar results such as in Syahrinudin (2005) where oil palms aged 3 and 10 years old in Sumatra, Indonesia resulted to a carbon stock of 9.15 tC/ha and 36.7 tC/ha, respectively which equates to an average of 22.92 tC/ha. Furthermore, in Yuliyanto et al (2016), oil palms aged 6-10 years old has a carbon stock equal to 31.86 tC/ha which coincides with the observed values for the same age range of oil palms in the study.

**Table 3.**
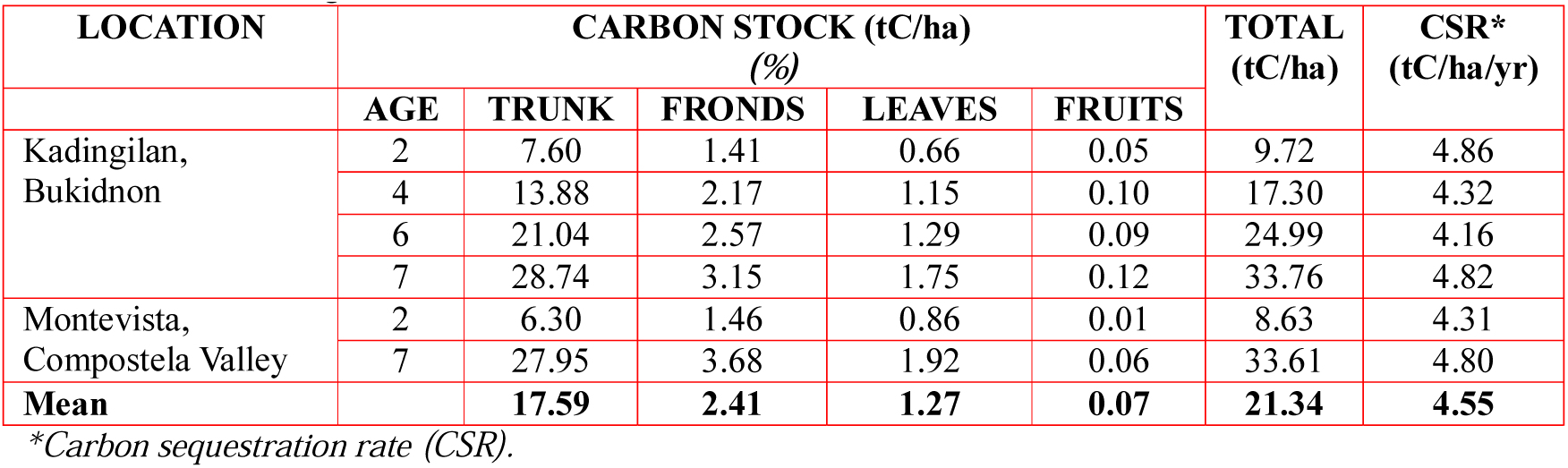
Carbon Storage in Oil Palm and its Parts

| LOCATION | CARBON STOCK (tC/ha) |  |  |  |  | TOTAL<br>(tC/ha) | CSR*<br>(tC/ha/yr) |
| --- | --- | --- | --- | --- | --- | --- | --- |
|  | AGE | TRUNK | FRONDS | LEAVES | FRUITS |  |  |
| Kadingilan,<br>Bukidnon | 2 | 7.60 | 1.41 | 0.66 | 0.05 | 9.72 | 4.86 |
|  | 4 | 13.88 | 2.17 | 1.15 | 0.10 | 17.30 | 4.32 |
|  | 6 | 21.04 | 2.57 | 1.29 | 0.09 | 24.99 | 4.16 |
|  | 7 | 28.74 | 3.15 | 1.75 | 0.12 | 33.76 | 4.82 |
| Montevista,<br>Compostela Valley | 2 | 6.30 | 1.46 | 0.86 | 0.01 | 8.63 | 4.31 |
|  | 7 | 27.95 | 3.68 | 1.92 | 0.06 | 33.61 | 4.80 |
| **Mean** |  | **17.59** | **2.41** | **1.27** | **0.07** | **21.34** | **4.55** |

\*Carbon sequestration rate (CSR).

Furthermore, oil palms in this study can be considered comparable to coconuts in terms of carbon stock. Carbon stored in coconuts has an average of 22.73 tC/ha based on a study in Sri Lanka (Ranasinghe and Thimothias 2012). However, bamboo plants (Patricio and Dumago 2014), fruit trees (Janiola and Marin 2016) as well as forest trees (Tulod 2015) in general have higher carbon stock than the oil palms in this study.

Moreover, majority of the carbon stock is found in the oil palm trunk which accounts for 73% to 84% of the total carbon stock. In the study by Pulhin et al (2014) most of the carbon in the oil palm plant is stored in the trunk.

In terms of the mean carbon sequestration rate, the average in the study is 4.55 tC/ha/year. This means that on the average, oil palms in this study can sequester this amount of carbon annually. Another study also has almost the same carbon sequestration rate of 4.9 tC/ha/year for a 10 old oil palm plantation in Indonesia (Syahrinudin 2005). However, a 9 year old oil palm plantation in the Philippines was estimated to be able to sequester 6.10 tC/ha/year (Pulhin et al 2014).

Previous studies show that the carbon sequestration rate of the oil palm plantations in this study is comparable to some forest plantations in the country (Lasco and Pulhin 2003) though majority of forest plantations may still have a higher carbon sequestration rate. Furthermore, a study in Sri Lanka suggests that oil palms can also have a comparable carbon sequestration rate with coconuts in low country dry zones in Sri Lanka with a rate of 4.8 tC/ha/year (Ranasinghe and Thimothias 2012). At the same time coconuts studied in Leyte, Philippines also has a carbon sequestration rate of 4.78 tC/ha/year (Lasco et al 2002).

### 3.3 Carbon Stock of the Different Carbon Pools in the Oil Palm Plantations

As shown in Table 4, the average carbon stock of oil palm plantations in the study is 40.33 tC/ha. A similar result (40 tC/ha) is reported in Sanquetta et al (2015) based on a study in an oil palm plantation in Brazil. Furthermore, considering only the aboveground biomass, a bit similar result was observed in Syahrinudin (2005) for a 10 year old palm plantation in Sumatra, Indonesia (38.94 tC/ha). Moreover, the above result are found to fall within the carbon stock values of coconut plantations in Sri Lanka (37 tC/ha – 64 tC/ha) (Ranasinghe and Thimothias 2012).

**Table 4.** Carbon Stock in the Different Carbon Pools of the Oil Palm Plantations

| LOCATION | CARBON STOCK (tC/ha) |  |  |  |  |
| --- | --- | --- | --- | --- | --- |
|  | PALM | UNDERSTORY | LITTER | SOIL | TOTAL |
| Kadingilan, Bukidnon | 21.44<br>(55%) | 1.41<br>(4%) | 2.34<br>(6%) | 13.99<br>(36%) | 39.18 |
| Montevista, Compostela Valley | 21.12<br>(51%) | 1.63<br>(4%) | 2.24<br>(5%) | 16.49<br>(40%) | 41.48 |
| **Average** | **21.28**<br>(53%) | **1.52**<br>(4%) | **2.29**<br>(6%) | **15.24**<br>(38%) | **40.33** |

*Values in parentheses are percentage distribution of carbon stock in each carbon pool.*

The above result however is considered low when compared to the carbon stock of fruit tree plantations (Janiola and Marin 2016), forest tree plantations (Tulod 2015), agroforestry farms (Labata et al 2012) as well as old growth forests (Lasco and Pulhin 2003). This means that conversion of forests to oil palm plantations will negatively impact the carbon balance of ecosystems. However, Sanquetta et al (2015) emphasized that permanent crops such as oil palm shows greater potential (in terms of carbon storage) when compared to temporary agricultural crops and livestock. In the same study it was cited that pasture land and agricultural lands only stores 8 tC/ha and 5 tC/ha, respectively. It is thus advisable that oil palm plantations serve as replacement for existing grassland as a strategy for the preservation of natural forest, avoiding emissions, as well as additional revenue (in the case of tradeable carbon) (Germer and Sauerborn 2008).

The majority of carbon stored in the plantations is in the oil palm which comprises 51% - 55% of the total carbon stock. Soil is the second major carbon pool which contains 36% - 40% of the total carbon stock. Understory and litter contain a combined average carbon stock percentage of 10%. This same trend is observed in forest plantations where trees generally contains majority of the carbon stock (Tulod 2015). This is contrary to the case of agroforestry farms wherein soil holds the most carbon (77-92%) (Labata et al 2012).

The average soil organic carbon in the oil palm plantations in this study is considered low (15.24 tC/ha) in comparison to other tree-based plantations (Labata 2012, Tulod 2015, Janiola and Marin 2016). In fact, such value is similar only to the soil organic carbon of pineapple plantations (14.72 tC/ha) and banana plantations (18.28 tC/ha) in Bukidnon, Philippines (Patricio 2014). By comparison, coconut plantations tend to have a higher soil organic carbon which is within 28.0 tC/ha – 30.3 tC/ha (Ranasinghe and Thimothias 2012; Patricio 2014).

Soil organic carbon is a significant carbon sink due to the fact that it is not released by burning and at the same time carbon remains in the soil for a longer time compared to other carbon pools (Lugo and Brown 1992). Previous studies have demonstrated the societal value of soil carbon emphasizing the need for its utilization, enhancement, and restoration by any means possible (Lal 2014, Siwar et al 2016). Related to that, it has been found out that while tree crop species improve biomass carbon, shrub crop species enhance soil carbon (Gnanavelrajah et al 2008). This could be the reason why oil palm plantations in the study have lesser soil carbon than biomass carbon (because oil palms can be equated to tree crop species in terms of ecosystem functions). Hence it would be beneficial to mix oil palm with shrub crop species in plantations to improve the carbon in the soil.

## 4. Conclusions

Oil palm plantations do have an essential role in climate change mitigation specifically in offsetting anthropogenic carbon emissions. This study shows that 2 to 7 years old oil palm plantations can store an average of 40.33 tC/ha and can sequester 4.55 tC/ha/year. Generally, oil palm plantations do not equal natural forests in terms of storing carbon, thus, it is not advisable to convert forest land into oil palm plantations. However, barren lands/grasslands are viable options because these lands have very low carbon stock. In fact, it has been previously shown that pasture land and agricultural lands have extremely lower carbon stock than oil palm plantations. Furthermore, based on the results, oil palm plantations are considered to have low soil organic carbon, an essential carbon pool in the ecosystem. Because shrub crop species are found to enhance soil organic carbon, it is recommended that these types of crops be incorporated with oil palms to complement the aboveground biomass in such plantations. With the present challenges regarding the streamlining of climate change mitigation in development planning, results of this study can provide basis for conceptualizing economic policies which are in harmony with environmental principles.

## Acknowledgments

The researchers would like to thank the owners of the oil palm plantations where this study was conducted. Likewise, the same gratitude is extended to the workers in the said plantations who were of great help especially during the data gathering stage of this research.

